# Proteome-wide cross-linking mass spectrometry to identify specific virus capsid-host interactions between tick-borne encephalitis virus and neuroblastoma cells

**DOI:** 10.1101/2021.10.29.464531

**Authors:** Sarah V. Barrass, Lauri I. A. Pulkkinen, Olli Vapalahti, Suvi H. Kuivanen, Maria Anastasina, Lotta Happonen, Sarah J. Butcher

**Author notes:** Corresponding authors (SB), (LH), (MA).

## Abstract

Virus-host protein-protein interactions are central to viral infection, but are challenging to identify and characterise, especially in complex systems involving intact viruses and cells. In this work, we demonstrate a proteome-wide approach to identify virus-host interactions using chemical cross-linking coupled with mass spectrometry. We adsorbed tick-borne encephalitis virus onto metabolically-stalled neuroblastoma cells, covalently cross-linked interacting virus-host proteins, and performed limited proteolysis to release primarily the surface-exposed proteins for identification by mass spectrometry. Using the intraviral protein cross-links as an internal control to assess cross-link confidence levels, we identified 22 high confidence unique intraviral cross-links and 59 high confidence unique virus-host protein-protein interactions. The identified host proteins were shown to interact with eight distinct sites on the outer surface of the virus. Notably, we identified an interaction between the substrate-binding domain of heat shock protein family A member 5, an entry receptor for four related flaviviruses, and the hinge region of the viral envelope protein. We also identified host proteins involved in endocytosis, cytoskeletal rearrangement, or located in the cytoskeleton, suggesting that entry mechanisms for tick-borne encephalitis virus could include both clathrin-mediated endocytosis and macropinocytosis. Additionally, cross-linking of the viral proteins showed that the capsid protein forms dimers within tick-borne encephalitis virus, as previously observed with purified C proteins for other flaviviruses. This method enables the identification and mapping of transient virus-host interactions, under near-physiological conditions, without the need for genetic manipulation.

**Author summary:** Tick-borne encephalitis virus is an important human pathogen that can cause severe infection often resulting in life-long neurological complications or even death. As with other viruses, it fully relies on the host cells, and any successful infection starts with interactions between the viral structural proteins and cellular surface proteins. Mapping these interactions is essential both for the fundamental understanding of viral entry mechanisms, and for guiding the design of new antiviral drugs and vaccines. Here, we stabilise the interactions between tick-borne encephalitis virus and human proteins by chemical cross-linking. We then detect the interactions using mass spectrometry and analyse the data to identify protein-protein complexes. We demonstrate that we can visualise the protein interaction interfaces by mapping the cross-linked sites onto the host and viral protein structures. We reveal that there are eight distinct sites on the outer surface of the viral envelope protein that interact with host. Using this approach, we mapped interactions between the tick-borne encephalitis virus envelope protein, and 59 host proteins, identifying a possible new virus receptor. These results highlight the potential of chemical cross-linking coupled with mass spectrometry to identify and map interactions between viral and host proteins.

## Introduction

Viruses are obligatory intracellular parasites that depend on virus-host protein-protein interactions (PPIs) to establish successful infections. The identification of these interactions and knowledge of the interaction interfaces contribute to our understanding of the initial steps of the viral life cycle, and can guide the design of antivirals and vaccines [1–6].

Advances in high throughput methods have led to the large-scale identification of virus-host interactions, but the structural characterisation of these interactions is often still limited [7]. Affinity purification coupled with mass spectrometry, yeast two-hybrid, and protein microarrays, have identified virus-host PPIs for multiple viruses including: Japanese encephalitis virus, H1N1 influenza, human immunodeficiency virus, human cytomegalovirus, and severe acute respiratory syndrome coronavirus 2 [8–12]. These methods are however limited in their applicability to detect transient interactions between wild-type viruses and cells. Alternative methods, using chemical cross-linking, or proximity labelling (BioID and TurboID), demonstrate improved detection of weak and transient interactions [13–18]. In these approaches, cells are probed with modified viral proteins conjugated to trifunctional cross-linkers or biotin ligases. Host proteins in close proximity to the viral bait are then permanently cross-linked or biotinylated, and purified using the biotin or cross-linker tag. Enriched proteins are detected by comparison of the protein signal to that in negative controls using bottom-up proteomics. Alternative chemical cross-linking mass spectrometry (XL-MS) workflows that directly detect the cross-linked peptides additionally provide information about the interaction interfaces. Previous studies have used the finite length of the chemical cross-linker to indicate the proximity of two amino acid side chains during the cross-linking reaction, and to build structural models of bacteria-host PPIs [19, 20]. This shows the potential of XL-MS in both the identification of PPIs and the characterisation of the binding interface.

The flavivirus, tick-borne encephalitis virus (TBEV) is the causative agent of one of the most important arbovirus-caused diseases in Europe, Russia, and Northern China [21, 22]. Symptomatic infection with TBEV can cause meningitis, encephalitis, and meningoencephalitis, and often results in life-long neurological complications or death [23, 24]. The TBEV virion has three different structural proteins, the envelope protein (E protein), membrane protein (M protein) and capsid protein (C protein), in addition to a lipid bilayer and an ∼11 kilobase-long positive-strand RNA genome (Fig 1). The E protein forms the smooth outer surface of the virion and is responsible for receptor binding [25, 26]. The atomic structure of the mature TBEV virion E and M proteins has been solved by cryo-electron microscopy at a resolution of 3.9 Å and the crystal structure of the E protein at a resolution of 1.9 Å (Fig 1) [25, 27]. Non-infectious immature and partially immature viruses also egress from cells, and have a spikey surface (Fig 1) [27]. No proteome-wide study of TBEV virus-host protein interactions has been published to our knowledge.

**Fig 1:**
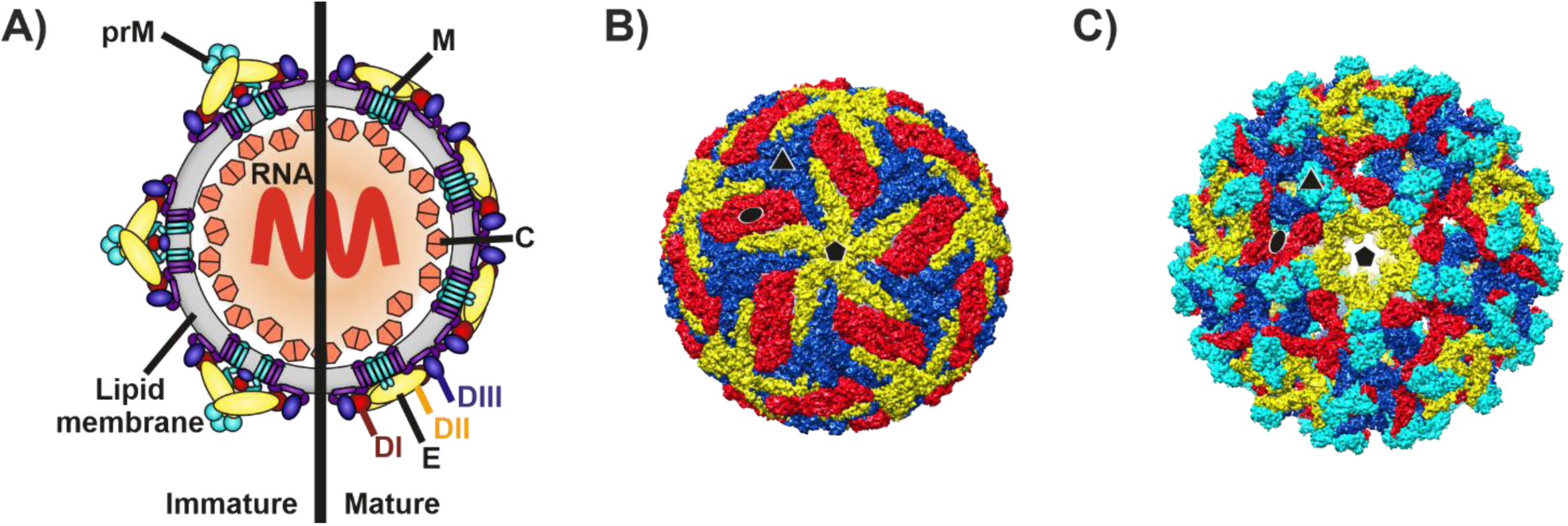
TBEV mature and immature virus structures. A) Schematic representation of TBEV with the immature structure shown on the left and the mature on the right. Multiple copies of the C protein dimer surround the genome forming the nucleocapsid complex, positioned beneath the lipid bilayer. The surface of the immature virion is covered in 60 spikes, composed of trimers of prM (pre-membrane)-E heterodimers embedded into the lipid bilayer. The surface of the mature virion is covered in 90 E-M heterotetramer complexes embedded into the lipid bilayer. TBEV E protein domain I is shown in red, domain III in dark blue, domain II in yellow and domain 4 in purple. B) Surface representation of the mature TBEV virion (PDB accession: 5O6A)[27]. The three E proteins within each asymmetric unit are shown in blue, red, and yellow. Symmetry axes are indicated by the black pentagon (five-fold), triangle (three-fold), and ellipse (two-fold). C) Surface representation of the immature Spondweni virus, a related flavivirus (PDB accession 6ZQW) [28]. The three E proteins within each asymmetric unit are shown in blue, red, and yellow, and the prM protein is shown in cyan. Symmetry axes are indicated by a black pentagon (five-fold), triangle (three-fold), and ellipse (two-fold).

In this large-scale proteomics study, we used XL-MS to identify the interaction interfaces of PPIs between TBEV and the surface of human neuroblastoma (SK-N-SH) cells. Here, the homobifunctional chemical cross-linker disuccinimidyl suberate (DSS) was used to covalently fix PPIs by cross-linking primary amine containing residues (the side chain of lysine residues or the N-terminus of the protein). The finite length of DSS (11.4 Å) imposed a maximum distance between cross-linked residues and was used to validate intraviral crosslinks by measuring their distances on TBEV proteins with known structures or reliable homology models. The final dataset was filtered using the intraviral cross-links as an internal control, leading to the identification 59 unique high confidence interactions between the TBEV E protein and cellular proteins.

## Results

### Identification of cross-linked peptides

To identify interactions between the mature TBEV virion and host proteins, we incubated the virus with metabolically-stalled neuroblastoma cells on ice, allowing for TBEV to bind to the cells, but preventing subsequent internalization. The TBEV-host PPIs were then stabilized and fixed by chemical cross-linking with DSS (Fig 2). To reduce the sample complexity and search space during data analysis, we used limited proteolysis to release primarily cell-surface associated host proteins. The released proteins were digested to peptides and analysed by liquid chromatography tandem mass spectrometry (LC-MS/MS) followed by label-free data dependent acquisition (DDA) quantitation to determine their relative abundance (S1 Table). Identified proteins were used to generate smaller, defined sets of sequences to use in the cross-linking data analysis workflow.

**Fig 2:**
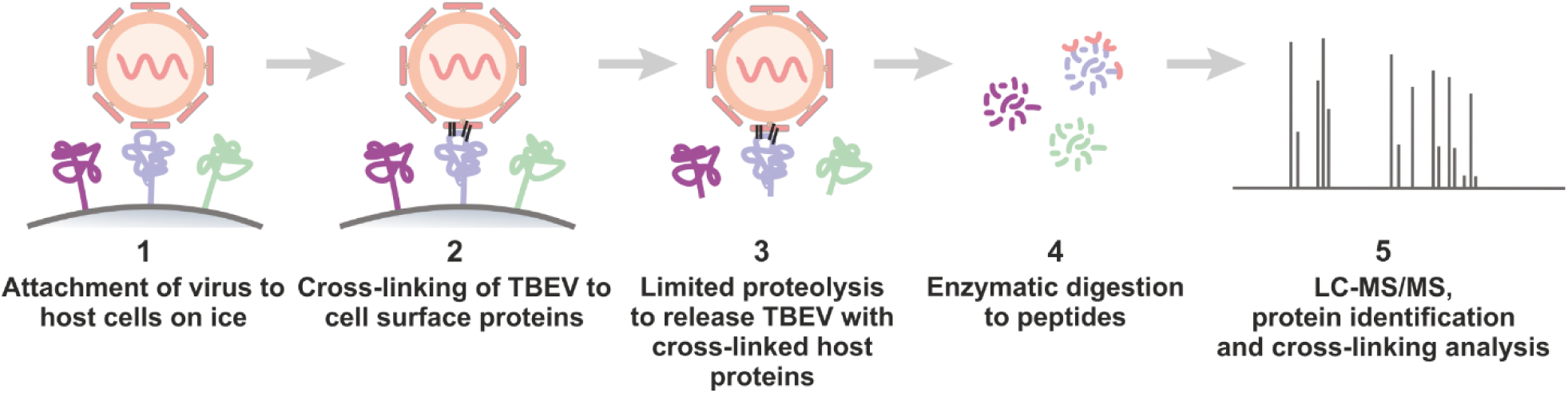
Schematic representation of cross-linking workflow. 1. TBEV was allowed to attach to metabolically-stalled SK-N-SH cells. 2. TBEV-host PPIs were stabilised by chemical cross-linking with DSS. 3. Cell-surface associated proteins, and cross-linked TBEV were released from the cell surface using limited proteolysis. 4. The released proteins were digested to peptides. 5. Peptides were analysed by LC-MS/MS and host and viral proteins identified and quantified using label-free DDA. Identified host proteins were analysed to identify cross-links between the host proteins and TBEV.

For cross-linking, we used four different cross-linker concentrations, in addition to a negative control sample to which no cross-linker was added. Each condition was repeated in triplicate, and the experiment repeated three times independently with different TBEV preparations and cell line passages, yielding 9 replicates per cross-linker concentration. The samples were initially analysed by immunoblotting of the TBEV E and C Proteins (Fig 3). The presence of higher molecular weight bands greater than 100 kDa in samples treated with DSS confirms cross-linking (Fig 3). The C protein has been shown to form antiparallel dimers in the crystal and NMR structures of other flaviviruses [29–31]. We identified a band with a molecular weight corresponding to that of C protein dimers, indicating that the C protein dimerizes in TBEV (Fig 3).

**Fig 3:**
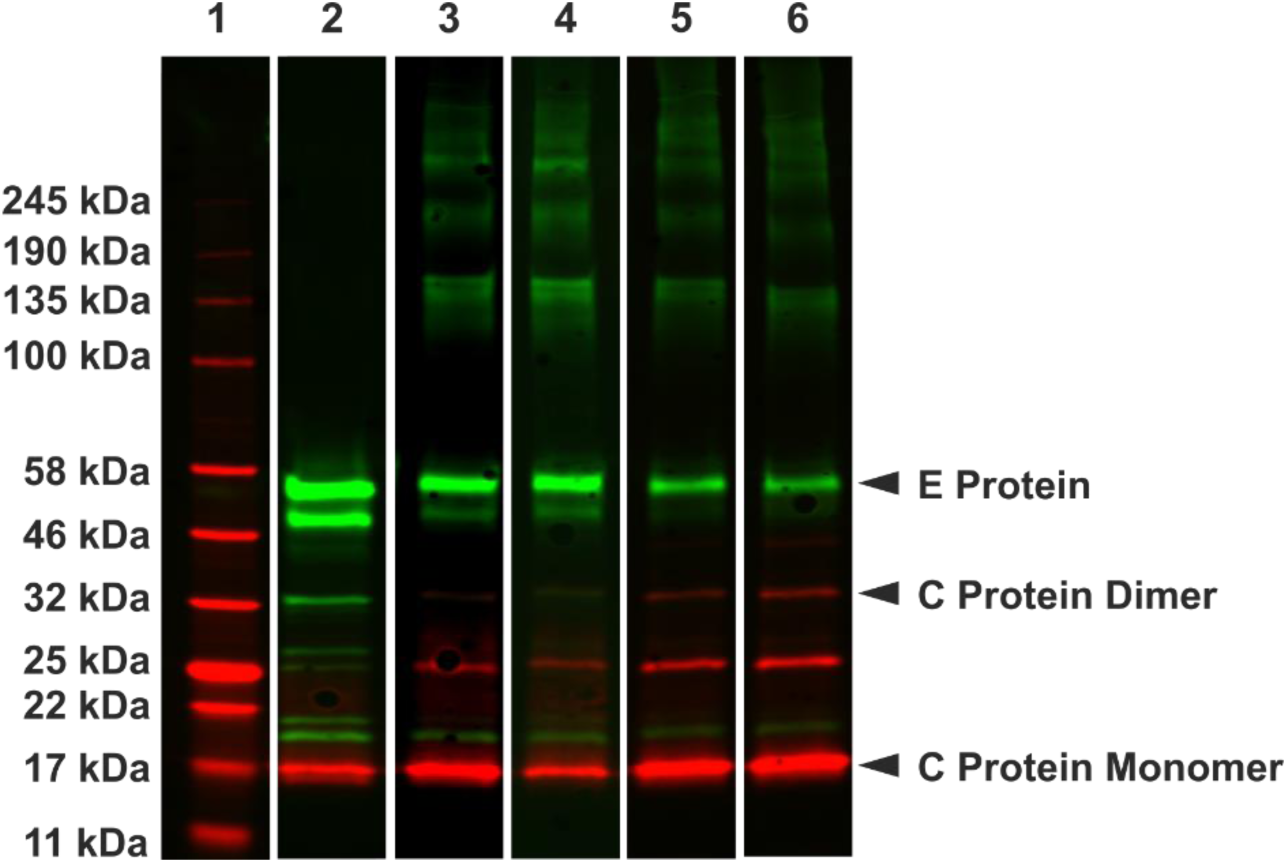
Immunoblot analysis of the TBEV E and C proteins in cross-linked samples. The TBEV C protein in shown in red and the TBEV E protein in green. Lane 1- protein marker; Lane 2- negative control with 0 mM DSS, Lane 3- cross-linking with 0.1 mM DSS; Lane 4- cross-linking with 0.25 mM DSS; Lane 5- cross-linking with 0.5 mM DSS; Lane 6- cross-linking with 1mM DSS. Higher molecular weight bands greater than 100 kDa corresponding to the cross-linking of E and C to other proteins can be seen in lanes 2-5. Lower molecular weight bands less than 50 kDa corresponding to the partial cleavage of the viral proteins during the limited proteolysis step are also observed in all lanes.

Identification of cross-linked peptides is computationally challenging as all primary amine-primary amine combinations in a given sequence database need to be considered. To reduce the search space for cross-linked peptide identification, proteins identified by DDA were probed for cross-links in batches (see materials and methods). A total of 7167 spectral observations of cross-linked peptides were identified using pLink2 at a false discovery rate (FDR) of 5%, excluding interfaces supported by cross-linked peptides identified in the negative controls, which correspond to false positives likely arising from erroneous peptide matches in the complex proteome background (S2 Table) [32]. Spectral observations of cross-links between two peptides within the same protein (intraprotein PPIs) accounted for the majority of the observations (5589), compared to 1578 for those identified between two peptides from different proteins (interprotein PPIs). Intraprotein cross-links have been consistently shown to make up a higher proportion of spectral observations within cross-linked datasets, as two residues within the same protein are highly likely to be in close physical proximity within the cell, leading to an increased cross-linking frequency [33, 34]. In total, 1697 different cross-linked interfaces were observed, and on average each interface was supported by 4.2 spectral observations. The cross-linked interfaces map to a network of 698 PPIs, consisting of 588 host proteins and the 3 viral structural proteins. Overall, 66.3 % of the unique PPIs were attributed to interactions between host proteins, 33.5 % to virus-host interactions and only 0.2 % to intraviral interactions. We also identified intraprotein cross-links in 297 host proteins and the TBEV C and E proteins.

The confidence of the cross-linking dataset can be investigated by examining the spatial distances between cross-linked residues for protein complexes where high-resolution structures or reliable homology models are available. As intraviral protein interactions account for 62 % of the detected spectral observations, we used the intraviral cross-links as an internal control to assess the confidence level for the dataset.

### Mapping of intraviral cross-links

The published mature TBEV structure and homology models of the immature virus and C protein dimer were used to accurately measure cross-link distances. The C-score of the homology models were -0.65 (C protein), 2.00 (E protein) and 0.57 (prM protein) [35–37] . We measured the distance between cross-linked residues and applied a maximum distance constraint of 30 Å between the lysine or N-terminus Cα (Table 1 and Figs 1 and 4) [38]. In total, 24 cross-links were mapped onto the viral structural proteins, and 22 fell within the accepted distance range. Overall, 14 of the cross-links satisfied the distance constraint in both the mature and immature TBEV structures, six were only acceptable in the mature structure and two in the immature structure. Cross-links with acceptable distances in only the mature structure were identified by 2089 spectral observations compared to 31 for those only accepted in the immature structure, demonstrating that the majority of the virus particles in the analysed samples were mature.

**Fig 4:**
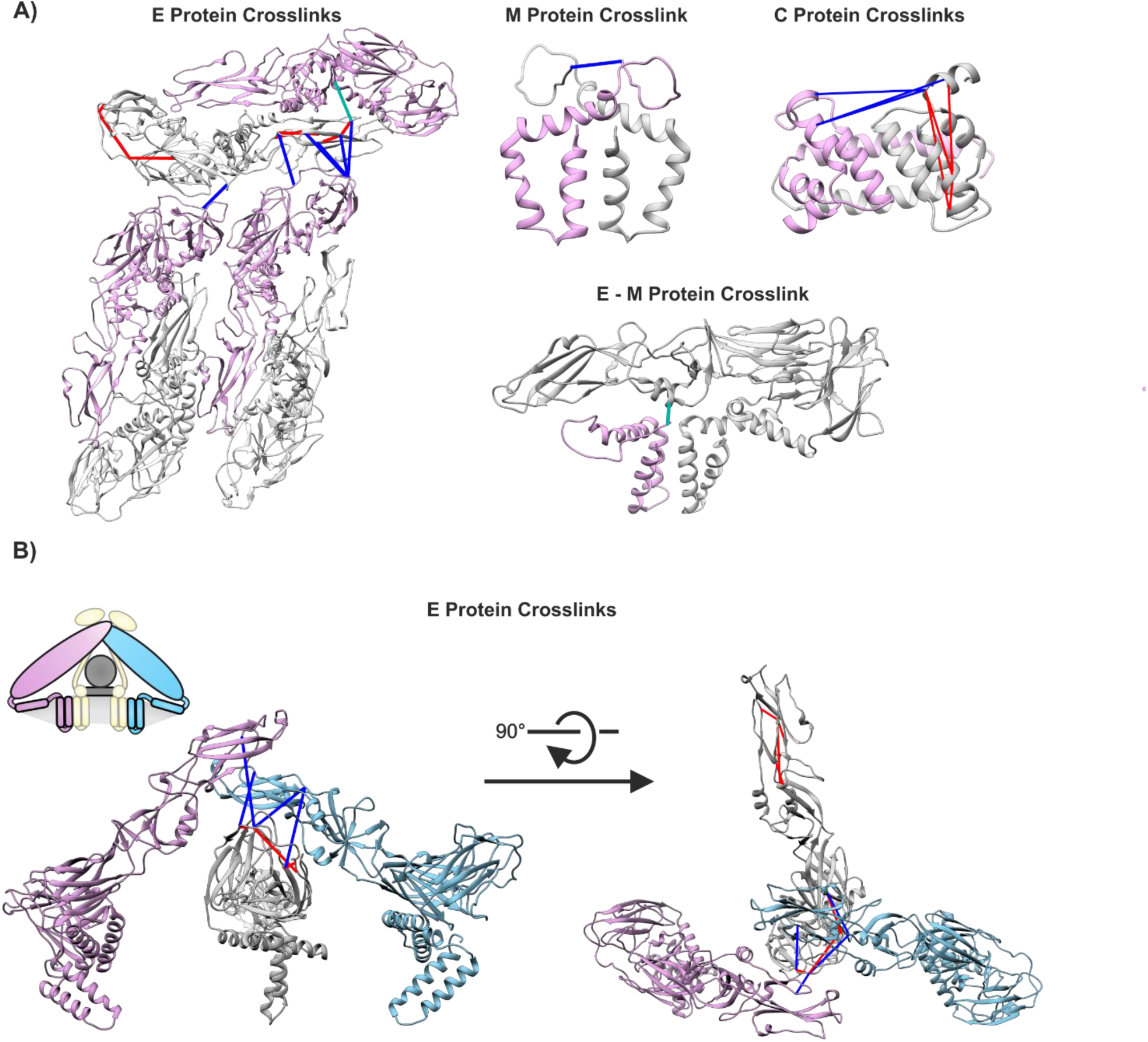
Mapping of intraviral crosslinks. Identified intraviral cross-links with the distances ≤ 30 Å mapped to the viral structural proteins are shown above in red, blue and teal. Red cross-links corresponding to intramonomer cross-links, teal intradimer cross-links, and blue interdimer or intertrimer cross-links. A) Cross-links mapped to the known cryoEM structure of the TBEV virion or the C protein homology model. B) Cross-links mapped to the generated homology model of the immature virus, E proteins in trimer 1 are coloured in plum and light blue and the E protein in trimer 2 is coloured grey. The schematic diagram additionally shows the position of the lipid bilayer (light grey) and the prM protein (light yellow).

**Table 1:**
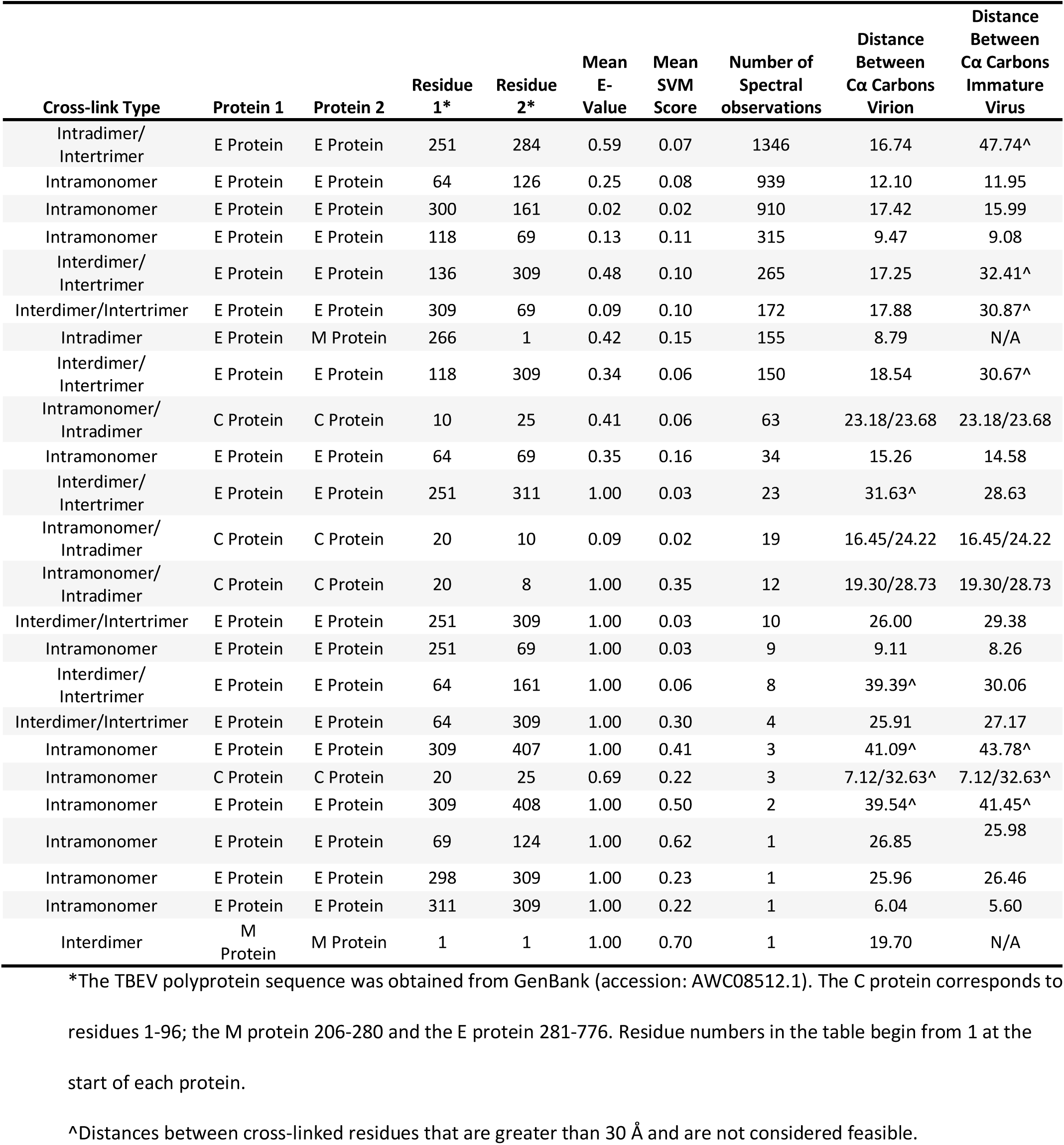
Intraviral cross-links identified, the corresponding average expectation value and SVM score of the peptide spectrum matches, and distances in the mature and immature virus structures

| Cross-link Type | Protein 1 | Protein 2 | Residue 1* | Residue 2* | Mean E-Value | Mean SVM Score | Number of Spectral observations | Distance Between Cα Carbons Virion | Distance Between Cα Carbons Immature Virus |
| --- | --- | --- | --- | --- | --- | --- | --- | --- | --- |
| Intradimer/Intertrimer | E Protein | E Protein | 251 | 284 | 0.59 | 0.07 | 1346 | 16.74 | 47.74^ |
| Intramonomer | E Protein | E Protein | 64 | 126 | 0.25 | 0.08 | 939 | 12.10 | 11.95 |
| Intramonomer | E Protein | E Protein | 300 | 161 | 0.02 | 0.02 | 910 | 17.42 | 15.99 |
| Intramonomer | E Protein | E Protein | 118 | 69 | 0.13 | 0.11 | 315 | 9.47 | 9.08 |
| Interdimer/Intertrimer | E Protein | E Protein | 136 | 309 | 0.48 | 0.10 | 265 | 17.25 | 32.41^ |
| Interdimer/Intertrimer | E Protein | E Protein | 309 | 69 | 0.09 | 0.10 | 172 | 17.88 | 30.87^ |
| Intradimer | E Protein | M Protein | 266 | 1 | 0.42 | 0.15 | 155 | 8.79 | N/A |
| Interdimer/Intertrimer | E Protein | E Protein | 118 | 309 | 0.34 | 0.06 | 150 | 18.54 | 30.67^ |
| Intramonomer/Intradimer | C Protein | C Protein | 10 | 25 | 0.41 | 0.06 | 63 | 23.18/23.68 | 23.18/23.68 |
| Intramonomer | E Protein | E Protein | 64 | 69 | 0.35 | 0.16 | 34 | 15.26 | 14.58 |
| Interdimer/Intertrimer | E Protein | E Protein | 251 | 311 | 1.00 | 0.03 | 23 | 31.63^ | 28.63 |
| Intramonomer/Intradimer | C Protein | C Protein | 20 | 10 | 0.09 | 0.02 | 19 | 16.45/24.22 | 16.45/24.22 |
| Intramonomer/Intradimer | C Protein | C Protein | 20 | 8 | 1.00 | 0.35 | 12 | 19.30/28.73 | 19.30/28.73 |
| Interdimer/Intertrimer | E Protein | E Protein | 251 | 309 | 1.00 | 0.03 | 10 | 26.00 | 29.38 |
| Intramonomer | E Protein | E Protein | 251 | 69 | 1.00 | 0.03 | 9 | 9.11 | 8.26 |
| Interdimer/Intertrimer | E Protein | E Protein | 64 | 161 | 1.00 | 0.06 | 8 | 39.39^ | 30.06 |
| Interdimer/Intertrimer | E Protein | E Protein | 64 | 309 | 1.00 | 0.30 | 4 | 25.91 | 27.17 |
| Intramonomer | E Protein | E Protein | 309 | 407 | 1.00 | 0.41 | 3 | 41.09^ | 43.78^ |
| Intramonomer | C Protein | C Protein | 20 | 25 | 0.69 | 0.22 | 3 | 7.12/32.63^ | 7.12/32.63^ |
| Intramonomer | E Protein | E Protein | 309 | 408 | 1.00 | 0.50 | 2 | 39.54^ | 41.45^ |
| Intramonomer | E Protein | E Protein | 69 | 124 | 1.00 | 0.62 | 1 | 26.85 | 25.98 |
| Intramonomer | E Protein | E Protein | 298 | 309 | 1.00 | 0.23 | 1 | 25.96 | 26.46 |
| Intramonomer | E Protein | E Protein | 311 | 309 | 1.00 | 0.22 | 1 | 6.04 | 5.60 |
| Interdimer | M Protein | M Protein | 1 | 1 | 1.00 | 0.70 | 1 | 19.70 | N/A |

\*The TBEV polyprotein sequence was obtained from GenBank (accession: AWC08512.1). The C protein corresponds to residues 1-96; the M protein 206-280 and the E protein 281-776. Residue numbers in the table begin from 1 at the start of each protein.
^Distances between cross-linked residues that are greater than 30 Å and are not considered feasible.

Locating the cross-links on the virus structures with distance constraints applied, allowed us to distinguish between three different types of E-E cross-links in the mature virion: intramonomer, intradimer and interdimer cross-links, and two in the immature virus: intramonomer and intertrimer cross-links. As expected, intramonomer E cross-links show similar distances in the mature and immature structures and accordingly the distance constraints are satisfied equally well for both. In contrast, as the E proteins rearrange from trimers to dimers upon virus maturation, only 2 cross-links satisfy the distance constraint in both the intertrimer positions found in the immature virus and the intradimer or interdimer positions found in the mature virus. Two types of cross-links are possible within the C protein dimer, intramonomer and intradimer. In our dataset, three of the four identified C protein cross-links satisfy the distance constraints for both interaction types, making it impossible for us to distinguish between these two alternatives.

Two of the intramonomer cross-links between E protein residues 309 and 407 or 408 demonstrate cross-linking lengths of ca. 40 Å in both the mature and immature structures. The cross-links were identified by a low number of spectral observations, 3 for 309-407 and 2 for 309-408, and may arise from interactions between disrupted virions, free E proteins, alternative viral conformations or be false positives. To distinguish between these alternatives, we examined the cross-linked spectra in detail (Fig 5). Fig 5A shows representative spectra for cross-links that satisfy the distance constraint and are identified by a high number of spectral observations, and Fig 5B and C show representative spectra for the two questioned crosslinks. As compared to the spectra shown in Fig 5A, spectra in Fig 5B and C show both a low signal-to-noise ratio and a low sequence coverage for the peptide fragments, and most likely represent false positive hits from background noise.

**Fig 5:**
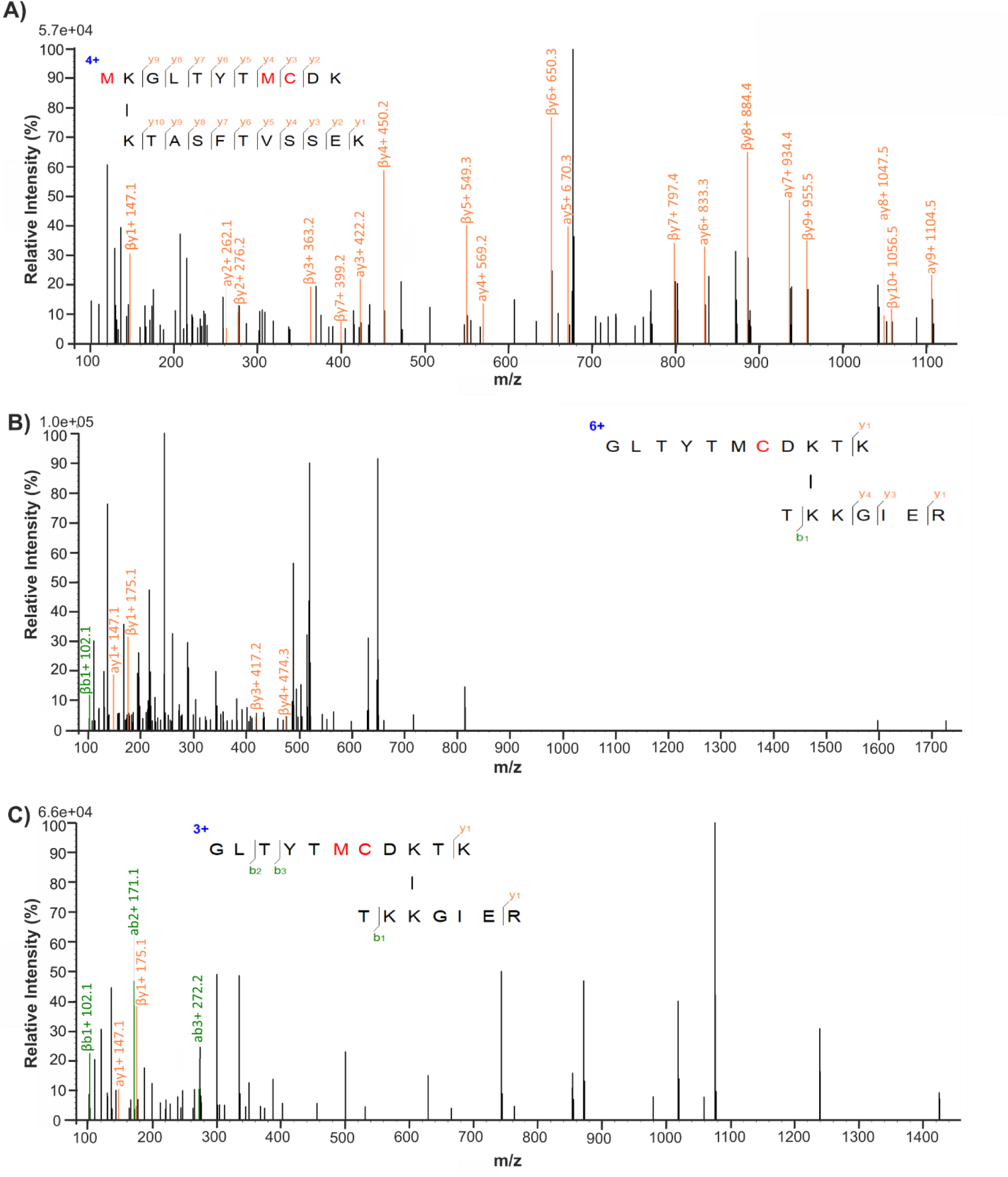
Representative spectra of E protein cross-links that satisfy the distance constraint (< 30 Å) and cross-links with distances > 30 Å. The cross-linked peptides are shown with the longer α peptide positioned above the shorter β peptide and a connecting line between the cross-linked residues. Modified residues in the peptides are shown in red and the detected beta and gamma fragments shown in green and orange respectively. Peaks corresponding to the detected fragments are coloured and labelled accordingly. The charge state of the cross-linked peptide is shown in blue. A) Representative spectrum of E protein cross-link 300-161 that are less than 30 Å and show a high number of spectral observations. B) Representative spectrum for the cross-link 309-407, that has a cross-linking distance of 41.09 Å in the mature structure and 43.78 Å in the immature structure. C) Representative spectrum for the cross-link 309-408 that has a cross-linking distance of 39.54 Å in the mature structure and 41.45 in the immature structure. Cross-links with distances > 30 Å show poor fragmentation and signal to noise ratio in the cross-linked spectral (B and C), compared to cross-links with distances < 30 Å (A).

Based on the imposed distance constraints, we calculated that 91.7 % of the intraviral cross-linked interfaces are identified with high confidence, at a distance threshold of <30 Å indicating that our approach allows detection of specific intra- and inter-protein cross-links with high confidence.

### Filtering the cross-linking dataset

The program pLink2 provides two parameters for assessing the confidence of cross-linked peptide spectrum matches (PSM), the expectation value (E-value) and the SVM score [32]. Both the E-value and SVM score describe the probability of a cross-linked PSM being a random match, and have values ranging from 1 to 0, where the smaller the value the more confident the PSM. The SVM score is calculated for every PSM, acting as the prime measure for FDR estimation, whereas the E-value is only calculated for PSMs that pass the FDR threshold [32]. In addition, cross-links may occur due to the sporadic proximity of proteins in the sample, or due to specific cross-linking of interacting proteins. Cross-links identified by more spectral observations have a higher probability of reflecting specific interactions. Therefore, the number of spectral observations per cross-linked interface can provide information about the cross-linking confidence on the protein-protein interaction level. Previous studies have used the E-value, SVM score, number of spectral observations or a combination of the aforementioned to assess the confidence of the identified cross-links but no standardized values have been established [32,39,40]. Here, we investigated the correlation between the E-value, SVM score, number of spectral observations and the acceptable distance constraint (<30 Å) as measured for our intraviral cross-links, to determine confidence parameters for the dataset.

Our data show a correlation between the measured distance of each unique intraviral cross-link and the calculated SVM scores, but not the E-values (Fig. 6, S1 Fig). Consequently, the E-value was not considered as a suitable measure of confidence for this dataset. We observed that intraviral cross-links with distances > 30 Å in both the mature and immature structures (indicated as red bars in Fig. 6) had the 3^rd^ and 4^th^ highest mean SVM scores of the intraviral cross-links (Table 1). Furthermore, cross-links with higher mean SVM scores were only identified by one spectral observation (Table 1). Based on the SVM scores of intraviral cross-links with distances > 30 Å in both the mature and immature structures, we imposed a SVM score threshold of < 0.355, hence discarding 884 of the identified cross-linked interfaces. Interestingly, 97.8 % of the excluded interfaces were identified by only one spectral observation.

**Fig 6:**
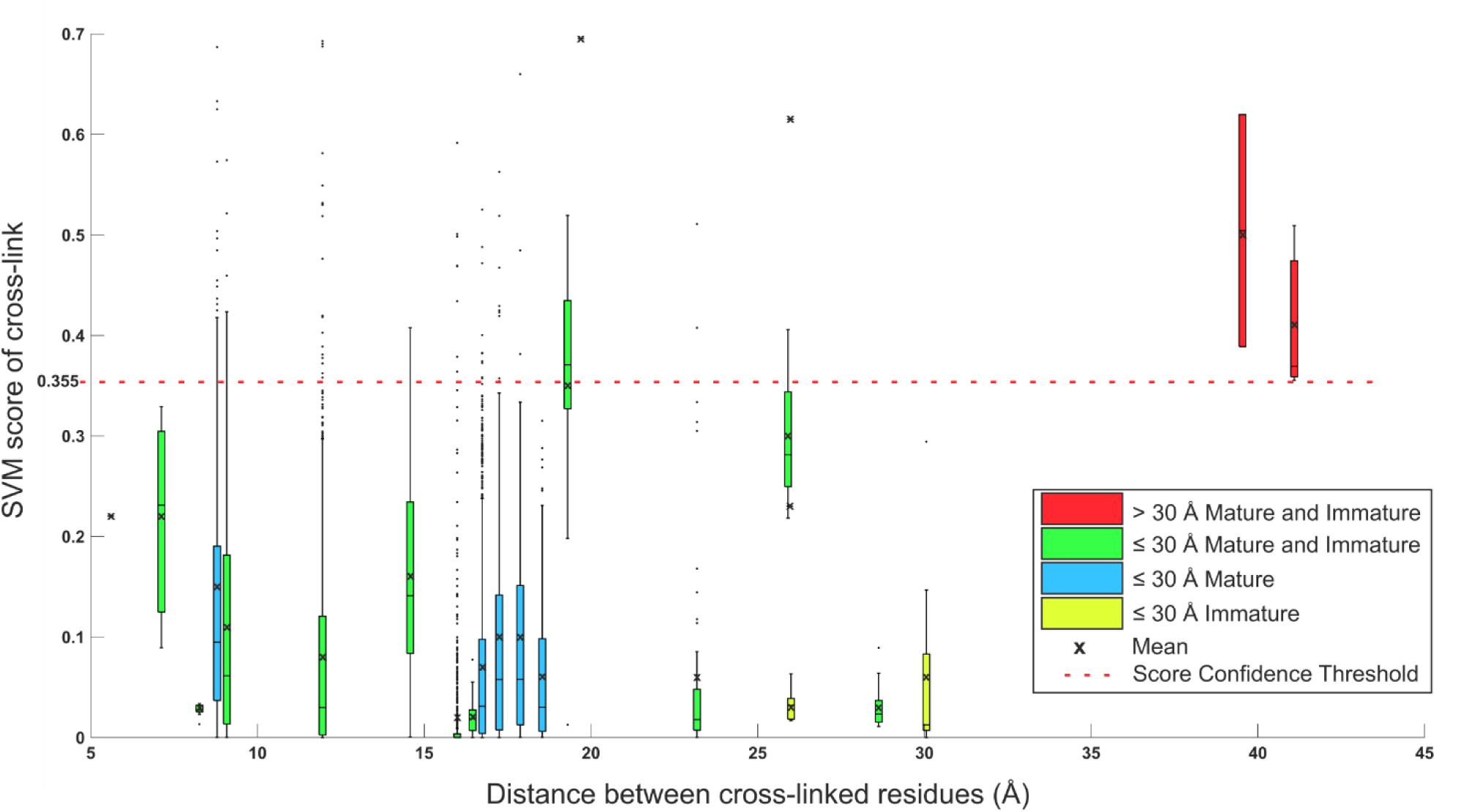
Box and whisker plot of the SVM scores for the intraviral cross-links, plotted against the distance between the cross-linked residues for each cross-link. The bars are coloured based on whether the distance constraint is satisfied for both the mature and immature conformations (green), only the mature conformation (blue), only the immature conformation (yellow) or not satisfied in either the mature or immature conformation (red). The mean SMV score is marked with a cross. A SVM score confidence threshold of < 0.355 is shown by a red dashed line.

The number of cross-linked spectral observations for 101 randomly selected proteins in the dataset was compared to the average spectral count of these proteins across all the samples in the DDA analyses (S1 File). The number of spectral observations for cross-linked peptide pairs for these same 101 selected proteins was also compared to the number of lysine residues present within each protein (S1 File). No correlation was observed in either of these cases; so, neither protein abundancy nor lysine content affect the data content (S1 File). Therefore, the number of spectral observations was used in a non-biased manner to assess the confidence of each unique cross-link to reduce the dataset for analysis further.

Here, we filtered the data in a stepwise manner, first on the PSM level using the SVM score and secondly on the protein-protein interaction level using the number of spectral observations. A cross-linked interface was considered to be of high confidence if it had an SVM score < 0.355 and was identified by ≥ 2 spectral observations. Filtering the data in the reverse order would lead to inclusion of interactions only supported by one high confidence spectral observation. The filtered cross-linking dataset consisted of 218 cross-linked interfaces, 36.7 % of which were attributed to virus-host PPIs (S3 Table).

### Virus-host protein-protein interactions

Virus-host PPIs were identified between TBEV and 61 host proteins in the filtered cross-linking dataset. In total, 59 proteins were shown to interact with the TBEV E protein and 2 with the M protein. Cross-links with the M protein form between N-terminal serine of mature M and the host proteins. As the M protein is buried and not accessible for cross-linking in the mature virion, these interactions likely occur between the host proteins and disrupted virions, free M protein, or conformations that have yet to be described.

E -host PPIs were mapped to eight different lysine residues on the outer surface of the virion (E protein residues 118, 126, 136, 161, 251, 280, 300, and 336; Fig 5), indicating that the host proteins were indeed interacting with assemble capsids. Only six of these residues shown in blue are also accessible on the outer surface of the immature particle, as residue 336 is obscured by other E proteins and residue 251 by prM (Fig 7). The cross-linked lysine residues are distributed evenly across the surface of the E protein, showing no preference for domains I, II or III (Fig 7).

**Fig 7:**
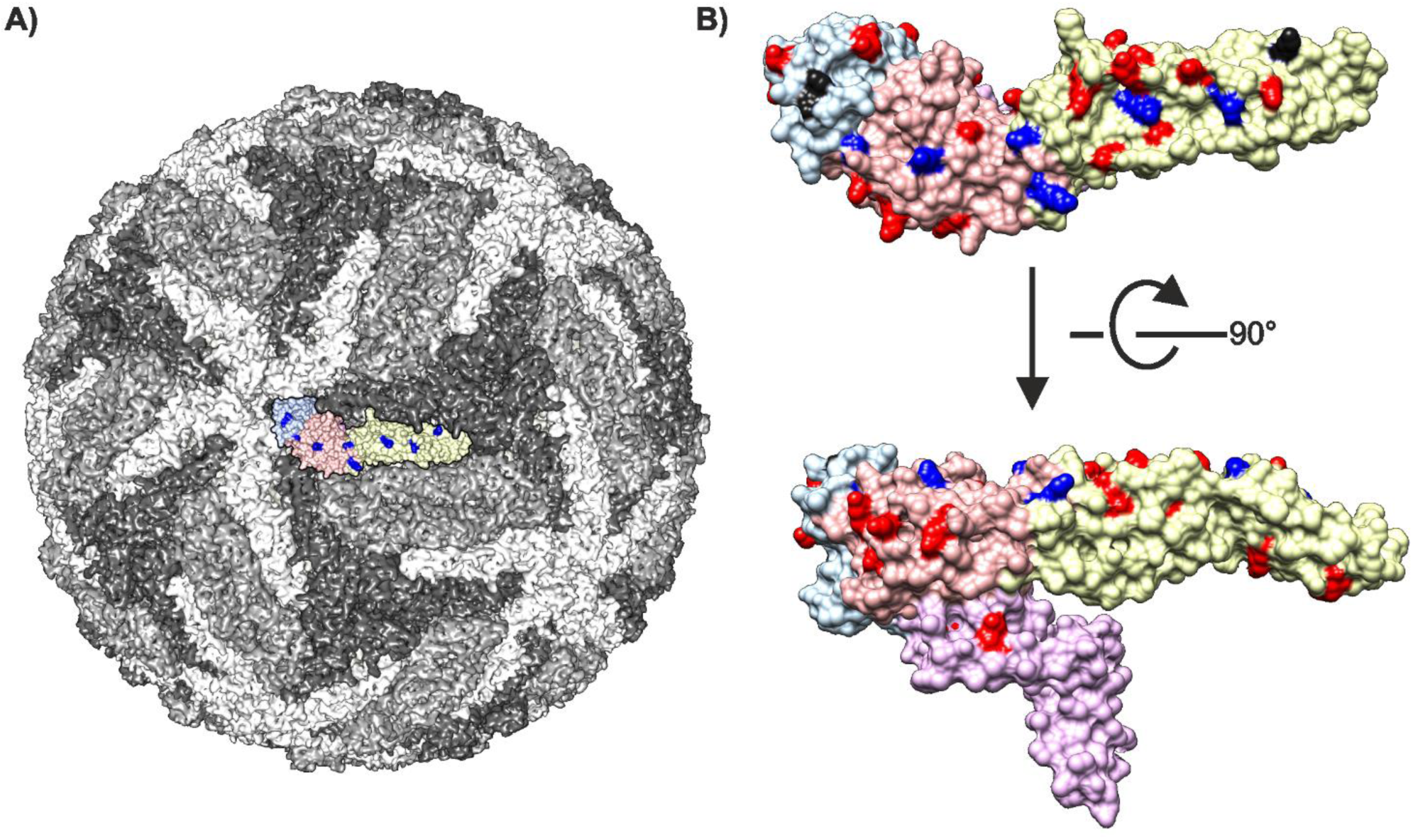
Visualisation of E protein lysine residues that form cross-links with host proteins. A) Surface representation of the TBEV virion (PDB accession: 5O6A) [27]. The three E proteins within each asymmetric unit are shown in white, grey and dark grey. One E protein monomer is shown in colour (TBEV domain I is shown in peach, domain II in yellow, domain III in lilac with the cross-linked lysines shown in blue. B) Mapping of the cross-linked lysines on the structure of the E protein (PDB accession 5O6A). Lysines detected with cross-links are shown in blue if located on the surface of both the mature and immature virus and black for those only located on the surface of the mature virus; lysines without cross-links are shown in red. TBEV domain I is shown in peach, domain II in yellow, domain III in light blue and domain IV in lilac.

**Fig 8:**
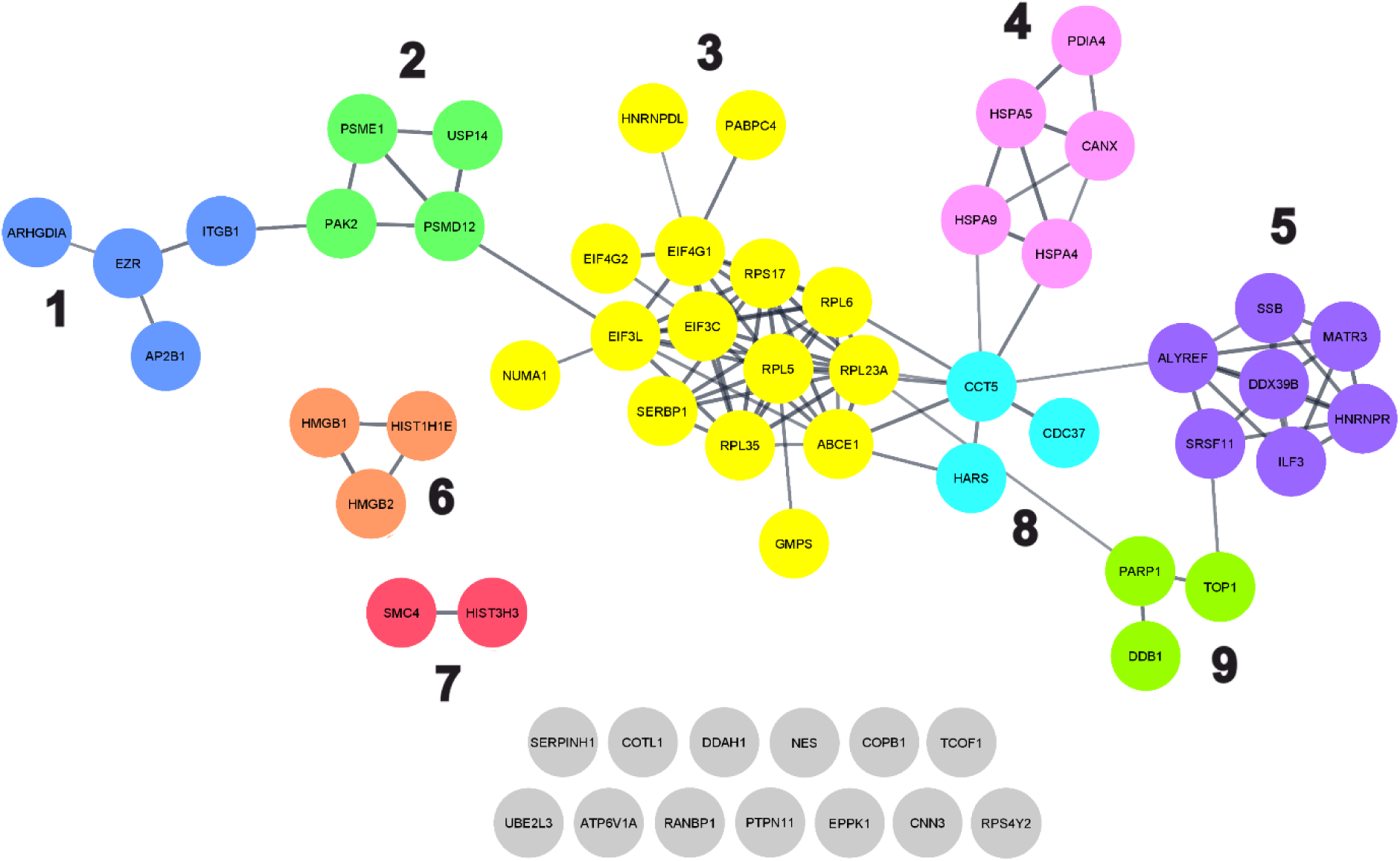
String network of 59 host proteins identified as interacting with the TBEV E protein in the filtered cross-linking dataset. The string-database network shows both direct and indirect protein interactions and is clustered based on the combined score associated with each interaction, using the Markov clustering algorithm [41]. Proteins are coloured based on the clustering and each cluster is numbered (1-9). Proteins that do not have any known interactions with other host proteins identified in the filtered cross-linking dataset are shown in grey.

We performed String-database analysis to identify if any of the 59 E protein-interacting host proteins also interact with each other (Fig 6) [41]. In total, 46 proteins were shown to interact with at least one other protein, and nine interaction clusters were identified. This suggests that TBEV may interact with both individual proteins and larger protein complexes. Gene ontology (GO) analysis (S4 Table) of the clusters indicates that proteins in clusters 1, 2 and 4 are located in the extracellular region (GO:0005576), at the plasma membrane (GO:0005886) or in extracellular exosomes (GO:0070062). Clusters 1 and 2 show enrichment for receptor-mediated biological processes, including receptor-mediated endocytosis (GO:0006898; cluster 1), and receptor-mediated signalling (GO:0038095, GO:0050852, GO:0002223; cluster 2), whereas proteins in cluster 4 are involved in protein transport (GO:0015031). Cluster 3 is the largest cluster and consists of proteins found in the cytosol (GO:0005829) predominantly as part of ribosomes (GO:0005840), ribonucleoprotein complexes (GO:1990904) or the eukaryotic translation initiation factor 4F complex (GO:0016281). Other cytosolic proteins belong to cluster 8 (GO:0005829) and are involved in protein folding (GO:0006457) or translation (GO:0006412). Finally, proteins in clusters 5, 6, 7 and 9 are primarily found in the nucleus (GO:0005634) and are involved in RNA processing (GO:0006396), posttranscriptional regulation of gene expression (GO:0010608; cluster 5), and chromosome organisation (GO:0051276; clusters 6, 7 and 9). Importantly, many proteins primarily located in the cytosol or nucleus (clusters 3, 5-9) are also found in the plasma membrane or extracellular regions where they perform alternative biological functions; for example, HMGB1 (UniprotKB:P09429; cluster 6), functions as a nonhistone nucleoprotein in the nucleus and an inflammatory cytokine in the extracellular region [42]. In addition, GO analysis identified 19 proteins that are associated with immune system process, and 11 proteins associated with the cytoskeleton or cytoskeletal rearrangement.

## Discussion

In this study, we present a chemical cross-linking proteomics approach to simultaneously identify TBEV- neuroblastoma cell PPIs and their interaction interfaces. We used metabolically-stalled cells to adsorb and cross-link virus only to the cell surface, hoping to primarily enrich proteinaceous cell surface interactions. The cross-linked proteins were released by limited proteolysis. Analysis of this highly complex protein mixture by LC-MS/MS generated a large database of spectra containing four different peptide species with the minority being cross-linked. In addition, cross-linked peptides are the least well-fragmented in the database. In the next step, the pLink2 software compares the database of spectra with all of the possible theoretical cross-linker reaction outcomes. This step is a clear bottle neck in the process as in our hands, only a subset proteins could be analysed at a time, requiring multiple batch runs. Here, we optimised the analysis workflow in order to extract the most significant virus-host PPIs from our complex data. Firstly, we reduced the cross-linking search space by only analysing proteins identified by linear peptides in the samples. Secondly, we have an internal validation control in the sample. We identified high-confidence intraviral crosslinks using both the known and predicted three-dimensional structures of TBEV and the known length of the chemical cross-linker. Then, we correlated the high confidence cross-links with the SVM score and the quality of the spectra, allowing us to use an SVM score cut-off < 0.355 for the entire dataset. Finally, we imposed a spectral count cut-off ≥ 2 to select for the most specific protein interactions. Using this method, we identified 22 high confidence unique intraviral cross-links and 59 high confidence unique virus-host PPIs between the surface of TBEV and human neuroblastoma cells. These proteins form a robust and reliable dataset that can be investigated further for their roles in the virus life cycle. They could be targets for intervention.

Our approach presents four major advantages over alternative approaches used to identify virus-host PPIs described in the literature: 1) The wild-type virus interacts with cellular proteins on the surface of neuroblastoma cells. In comparison, affinity purification, yeast two hybrid and protein microarrays detect interactions in artificial systems [8–12]. Therefore, the expression levels, presentation and glycosylation state of the host and viral proteins may differ from that in natural infections, leading to both false positives and false negatives; 2) When fully assembled viral particles are used as bait, the capsid proteins are in the right biological conformation and molecular context for infection. In contrast, single recombinant bait proteins used in affinity purification, yeast two hybrid, protein microarrays and BioID may not be [8–12,16]. For instance, the monomeric Dengue virus ancillary receptor, DC-SIGN binds across two neighbouring E proteins on the capsid [43]. In our virus-based protocol, there would be 90 such sites for the DC-SIGN interaction giving both the correct biological context, but also increasing the avidity of the interactions; 3) The virus interacts with cellular proteins prior to crosslinking. In contrast, in the previously described approaches using trifunctional cross-linkers or BioID, the viral proteins are first conjugated to the cross-linker or biotin ligase prior to the interaction with cells. Conjugation may require genetic modification [13–17]. Consequently, interactions may be missed if the modification sterically hinders the binding region. Here we could identify PPIs, under near-physiological conditions, without the need for genetic manipulation; 4) The clearest advantage in our approach as compared to other methods mapping virus host PPIs in culture is the use of DSS. Despite its limitations (see below), this allows us to directly map peptide-level interaction interfaces. Other approaches lacking this level of information have been used for instance for identifying host receptors to Sars-CoV using crosslinking followed by immunoprecipitation and LC-MS/MS from SDS-PAGE bands, identifying vimentin as a critical protein for virus entry [18]; or using an affinity-enrichable crosslinker to identify NCMA1 as a receptor for Zika virus [15].

Although this approach shows promise for detecting a wide-range of virus-host PPIs, there are also some challenges. The complete coverage of the interaction space is limited by the accessibility of surface cross-linkable lysine residues and the sample complexity. In order to detect an interaction, there must be cross-linkable residues on both sides of the interaction interface within 30 Å of each other. Furthermore, in complex systems containing a higher number of protein species, the number of cross-links per species is lower in comparison to simpler systems. Consequently, interactions that occur in lysine deficient regions or with low frequency cannot be detected, leading to an incomplete picture of the interaction interface or failure to detect the PPI. Performing parallel experiments using chemical cross-linkers with different lengths or reactive residues such as arginine, aspartate or glutamate, could overcome this limitation [44, 45]. Simple cross-linking experiments including only the virus and a single interaction partner can be used to ensure complete coverage of the 3D interaction space once interesting PPI have been identified.

Having considered the potential advantages and challenges of this protocol, we will now consider potential biological implications. Laminin binding protein has previously been suggested as a TBEV receptor. It was present in our dataset, but no cross-links were identified to TBEV [46, 47]. Our data do not support that laminin binding protein is a TBEV receptor in this cell line. However, we have identified proteins that are associated with the early stages of viral infection in other viruses, including ITGB1, ATP6V1A, EZR, HSPA9, and HSPA5. ITGB1 as an entry receptor for a large number of viruses including, cytomegalovirus, Epstein-Barr virus, human parvovirus B19, and mammalian reovirus [48–51]. ATP6V1A directly interacts with rabies viral matrix protein facilitating uncoating [52]. EZR is an essential host factor required for the entry of Japanese encephalitis virus into human brain microvascular endothelial cells [53]. HSPA9 has been identified as a putative receptor for Tembusu virus [54].

HSPA5 (Cluster 4, Fig 6) has been identified as a receptor for several flaviviruses including, Zika and Japanese encephalitis where HSPA5 was shown to affinity purify with recombinant E protein domain III, but the interaction context in virions has not been studied [56, 57]. HSPA5 is a multifunctional regulator of endoplasmic reticulum homeostasis, playing an important role in protein processing and quality control [59–62]. It is located both in the endoplasmic reticulum lumen and on the outer surface of the plasma membrane in many cell types including neurons [63–66]. HSPA5 consists of 2 domains, a nucleotide-binding domain that binds ATP, and a substrate-binding domain that binds and stabilises partially folded or folded proteins [67]. Here we identified four spectral observations mapping to the TBEV E hinge region (residue 136) and the HSPA5 substrate binding domain residues 521 and 516 (Fig 9). Therefore, the interaction between HSPA5 and the E protein hinge region could constitute a unique binding interface, or be part of a larger interface that also binds domain III consistent with other flavivirus studies [54–56,58]. We hypothesize that HSPA5 could interact with both the hinge region of one E monomer and domain III of an adjacent E monomer at the 3-fold axis, where the regions are in close spatial proximity (Fig 9). Interestingly, Fab fragments of the TBEV neutralising antibody 19/1786 have been shown to bind across this interface at the 3-fold axis potentially preventing the HSPA5 interaction[27]. Although, no cross-links were detected between domain III and HSPA5 in our study, this can be partly explained by poor lysine availability. Structural bioinformatics studies of the Zika virus domain III-HSPA5 interaction do not detect any interacting lysine residues on HSPA5[68].

**Fig 9:**
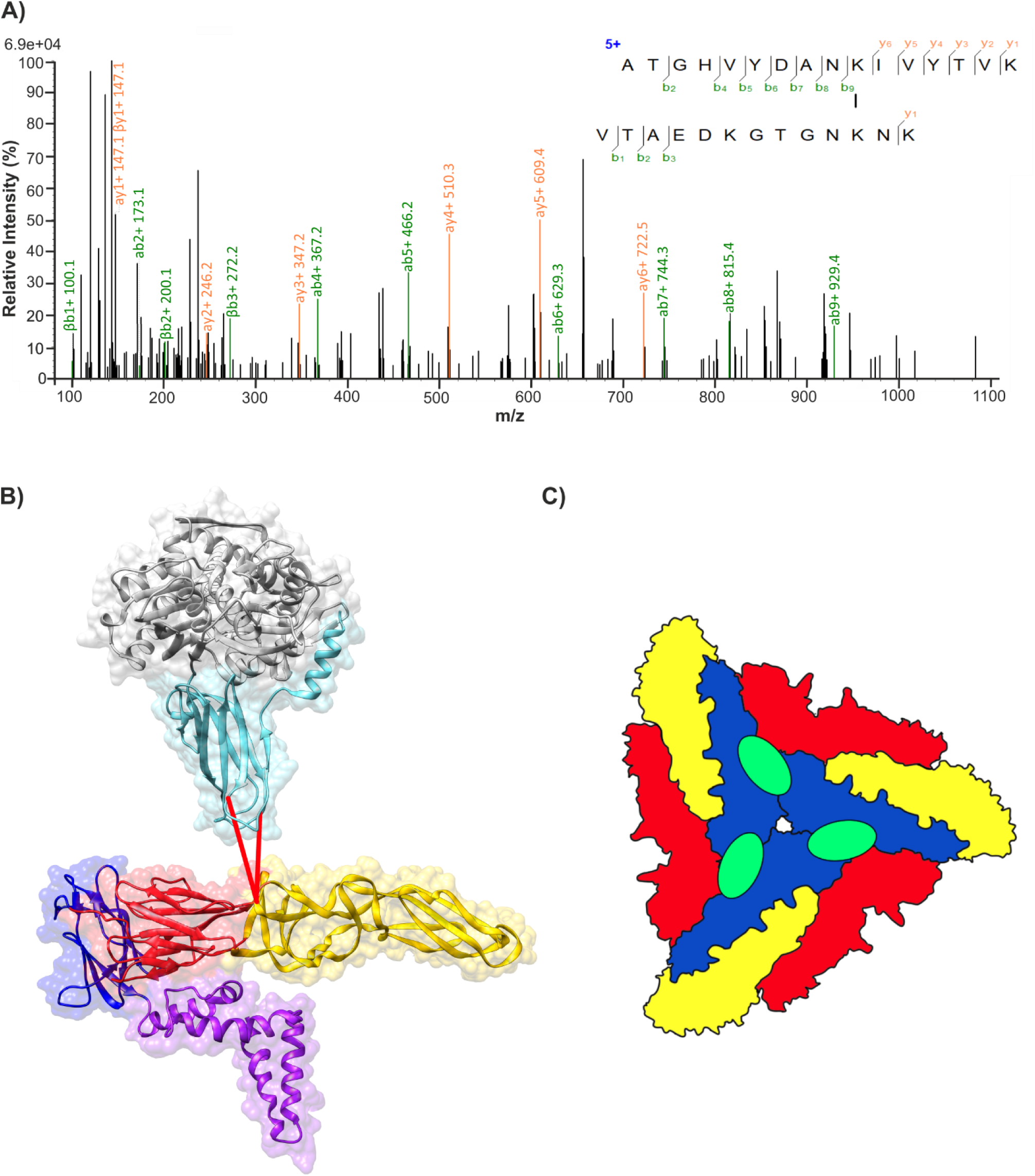
Cross-linking of HSPA5 and TBEV E protein. A) Assigned spectrum for the peptide associated with the HSPA5-E protein interaction. B) TBEV E protein (PDB accession 5O6A) and HSPA5 (PBD accession 6ZMD) were placed in close proximity to allow for the visualisation of the cross-links, indicated here with red lines. TBEV E protein domain I is shown in red, domain III in dark blue, domain II in yellow and domain 4 in purple. HSPA5 substrate-binding domain is shown in light blue and the remainder of the protein in grey. C) Schematic representation of speculated HSPA5 binding region on the mature virus. The three E proteins within each asymmetric unit are shown in blue, red, and yellow. The proposed HSPA5 binding site is shown in green.

After the virus has attached to the cell surface, the next step is to enter the cells either through clathrin-mediated endocytosis or macropinocytosis, and we found evidence for both pathways being used. We identified AP2B1 in the GO analysis that binds to the clathrin heavy chain [69]. We also identified an abundance of cytoskeletal proteins and cytoskeletal remodelling proteins required in macropinocytosis including, NES, NUMA1, ITGB1, EZR, PAK2, COTL1, CCT5, CNN3, EPPK1 and RANBP1. This supports the hypothesis that macropinocytosis is used by TBEV as well as Dengue virus [70].

## Conclusions

In this study, we present a XL-MS method to identify and map transient virus-host PPIs under near-physiological conditions, without the need for genetic modification. Using this method, we identified 59 high confidence virus-host PPIs between TBEV and the surface of neuroblastoma cells. These proteins form a robust and reliable dataset that can be investigated further for functional relevance in targeted follow-up experiments. The presented methodologies are generally applicable to other virus-host systems and can assist in expanding our knowledge of viral infections.

## Materials and methods

### Viruses and cells

Neuroblastoma SK-N-SH cells (ECACC 86012802) were maintained in Dulbecco’s modified Eagle’s medium (DMEM) (Sigma) supplemented with 10 % heat inactivated FBS (Gibco), 100 μg/ml of penicillin-streptomycin mix (PenStrep) (Sigma) and 2 mM L-glutamine (Sigma), at 37 °C and under a 5 % CO_2_ atmosphere. For virus propagation, cells were infected with TBEV strain, MG569938 Kuutsalo-14 *Ixodes ricinus* Finland-2017, at a MOI of 0.003, in Dulbecco’s modified Eagle’s medium (DMEM) (Sigma) supplemented with 2 % heat inactivated FBS (Gibco), 100 μg/ml of penicillin-streptomycin mix (PenStrep) (Sigma), 2 mM L-glutamine (Sigma) and 0.35 µM rapamycin and incubated at 37 °C for 3 days under a 5 % CO_2_ atmosphere. Virus-containing supernatant was aspirated and centrifuged at 4500 x g for 5 minutes to remove cell debris. Cleared supernatant was aliquoted into cryopreservation vials and frozen at -80 °C. Viral titers were determined using plaque-forming assay. SK-N-SH cells were grown on a 6-well plate, infected with ten-fold serial dilutions of the virus and incubated for 1 hour at 37 °C, under a 5 % CO_2_ atmosphere. Overlay medium (Minimum Essential Medium Eagle, PenStrep and 2 mM L-glutamine, 1.2 % avicel) was added to each well and the cells incubated for 4 days at 37 °C, under a 5 % CO_2_ atmosphere. After 4 days the cells were fixed with 10 % formaldehyde, stained with crystal violet and the plaques counted to determine the number of plaque forming units per ml (PFU/ml).

### Production of amino acid free virus stock

SK-N-SH cells were grown to 90% confluence, washed twice with PBS and the medium changed to amino acid free DMEM (Genaxxon) supplemented with 100 μg/ml of penicillin-streptomycin mix (PenStrep) (Sigma) and rapamycin 0.35 µM (selleckchem). Cells were infected with TBEV, at multiplicity of infection 1, and incubated at 37 °C for 4 Days, under a 5 % CO_2_ atmosphere. Virus-containing supernatant was aspirated and centrifuged at 4500 x g for 5 minutes to remove cell debris. The virus was pelleted through a 30 % sucrose cushion in HNE buffer (20 mM HEPES pH 8.5, 150 mM NaCl, 1 mM EDTA), 2 h, 27000 rpm, 4 °C. The virus pellet was then resuspended in amino acid free DMEM (Genaxxon) overnight, 4 °C, with mild shaking. The virus was aliquoted into cryopreservation vials and frozen at -80 °C. The stock was titered as described above. The typical obtained titer was 5 x 10^9^ pfu/ml.

### Cross-linking of TBEV with SK-N-SH cells

SK-N-SH cells were grown in a 6 well plate to 90 % confluence, washed twice with PBS, and the medium changed to amino acid free DMEM (Genaxxon) supplemented with 100 μg/ml of penicillin-streptomycin mix (PenStrep) (Sigma). The cells were incubated overnight at 37 °C, under a 5 % CO_2_ atmosphere. The cells were infected with TBEV at a MOI of 375 for 60 min, with rocking, on ice. Heavy/light disuccinimidyl suberate cross-linker (DSS-H12/D12, Creative Molecules Inc., www. creativemolecules.com) resuspended in dimethylformamide (DMF) was added to final concentrations of 0, 100, 250, 500 and 1000 µM and incubated for 60 min, with rocking on ice. The cross-linking reaction was quenched with a final concentration of 50 mM ammonium bicarbonate with rocking, on ice. The SK-N-SH cell surface proteins with attached TBEV virions were digested off with 1.25 μg trypsin (Promega) and the supernatant collected. Finally, cell debris was removed via centrifugation (16,000 x g, 5 min), the supernatant recovered, and the samples prepared for mass spectrometry (Fig 2).

### Immunoblot analysis

Proteins were resolved in 4-20 % SDS-PAGE, transferred onto a nitrocellulose membrane, and probed using anti-Langat E protein (BEI NR-40318; 1:1000 dilution) and C protein (57) (1:1000 dilution) antibodies in 5 % milk, tris-buffered saline 0.1 % tween-20 (TBST) [71]. The protein bands were visualised using IR800 and IR680-conjugated secondary anti-rabbit (Li-COR, 926-68071) and anti-mouse antibodies (KPL, 072071806) diluted 1:10,000 in tris-buffered saline 0.1 % tween-20 (TBST). The membrane was imaged using the Odyssey infrared imaging system (Li-COR).

### Preparation of cross-linked samples for mass spectrometry

Samples from cross-linking were first denatured with 8 M urea-100 mM ammonium bicarbonate. The cysteine bonds were then reduced with 5 mM tris(2-carboxyethyl) phosphine (37 °C, 60 min, 400 rpm) and alkylated with 10 mM 2-iodoacetamide (22 °C, 30 min, in the dark). Protein digestion was then performed with 0.1 µg/µl sequencing-grade lysyl endopeptidase (Wako chemicals) (37 °C, 2h, 400 rpm). Following the dilution of the sample with 100 mM ammonium bicarbonate to a final urea concentration of 800 mM the proteins were digested further with 0.2 µg/µl trypsin (Promega) (37 °C, 18h, 400 rpm). Digested samples were then acidified with 10% formic acid to a pH of 3.0, and the peptides were subsequently purified with C18 reverse-phase spin columns according to the manufactures protocol (Microspin Column, SS18V, The Nest Group, Inc). Peptides were then dried in a speedvac and reconstituted in 2% acetonitrile, 0.2% formic acid prior to mass spectrometric analyses

### Liquid chromatography tandem mass spectrometry

All peptide analyses were performed on a Q Exactive HFX mass spectrometer (Thermo Scientific) connected to an EASY-nLC 1200 ultra-high-performance liquid chromatography system (Thermo Scientific). The peptides were loaded onto an Acclaim PepMap 100 (ID 75μm x 2 cm, 3 μm, 100 Å) pre-column and separated on an EASY-Spray column (Thermo Scientific; ID 75 μm × 25 cm, column temperature 45 °C) operated at a constant pressure of 800 bar. A linear gradient from 4% to 45% of 0.1% formic acid in 80% acetonitrile was run for 50 min at a flow rate of 300 nl/min One full MS scan (resolution 60,000@200 m/z; mass range 350 to 1600 m/z) was followed by MS/MS scans (resolution 15,000@200 m/z) of the 15 most abundant ion signals. The precursor ions were isolated with 2 m/z isolation width and fragmented using higher-energy collisional-induced dissociation at a normalized collision energy of 30. Charge state screening was enabled, and precursors with an unknown charge state and singly charged ions were excluded. The dynamic exclusion window was set to 15 s and limited to 300 entries. The automatic gain control was set to 3 x 10^6^ for MS and 1 x 10^5^ for MS/MS with ion accumulation times of 110 and 60 ms, respectively. The intensity threshold for precursor ion selection was set to 1.7 x10^4^.

### MS data analysis

Raw DDA data was converted to gzipped and Numpressed mzML [72] using MSconvert from the ProteoWizard, v3.0.5930 suite [73]. All data was managed and analysed using openBIS [74]. The acquired spectra were analysed using the search engine X! Tandem (2013.06.15.1-LabKey, Insilicos, ISB) [75], OMSSA (version 2.1.8) [76] and COMET (version 2014.02 rev.2) [77] against an in-house compiled database containing the reviewed *Homo sapiens* reference proteome (UniProt proteome ID UP000005640) and the TBEV proteome (GenBank accession: AWC08512.1), yielding a total of 78121 protein entries and an equal amount of reverse decoy sequences. Full tryptic digestion was used allowing two missed cleavages. Carbamidomethylation (C) was set to static and oxidation (M) to variable modifications, respectively. Mass tolerance for precursor ions was set to 0.2 Da, and for fragment ions to 0.02 Da. Identified peptides were processed and analysed through the Trans-Proteomic Pipeline (TPP v4.7 POLAR VORTEX rev 0, Build 201403121010) using PeptideProphet [78]. The false discovery rate (FDR) was estimated with Mayu (v1.7)[79] and peptide spectrum matches (PSMs) were filtered with protein FDR set to 1% resulting in a peptide FDR >1%. Proteins were filtered to remove hits identified by only 1 unique peptide, and an average spectral count of < 2 across all samples. The protein names and corresponding UniProt ID are given in Supplementary information 1.

### Cross-link identification

In total 874 proteins that were identified by 2 or more unique peptides, and an average spectral count of ≥2 across all samples in the DDA data were probed for cross-links. Cross-links between the TBEV structural proteins and host proteins were identified using the pLink2 software package [32]. In order to reduce the search space for cross-link identification the data was analysed in 35 batches. The raw data dependent acquisition data, and compiled FASTA file databases containing a total of 28 protein sequences, from 25 different host proteins and the 3 viral structural protein were used as the software input. Host protein sequences were obtained from UniProt. The TBEV polyprotein sequence was obtained from GenBank (accession: AWC08512.1), the C protein corresponds to residues 1-96, the M protein 206-280 and the E protein 281-776. The following search parameters were used in the pLink2 software: Conventional cross-linking (Higher-energy C-trap dissociation (HCD)), precursor mass tolerance of 20 ppm; fragment mass tolerance of 20 ppm; peptide length of 6-60; peptide mass of 350-6000 Da, up to 3 missed cleavage sites; carbamidomethylation (C) was set to static modification; and oxidation (M) to variable modification. The results were then filtered using a filtering tolerance of ±10 ppm and a separate FDR >5% at the peptide spectrum matches level. 7716 cross-linked spectral observations were observed across all samples, 302 of these were observed in the negative control samples. In total, 247 cross-linked spectral observations in the 0.1-1mM DSS samples corresponded to cross-linked interfaces also identified in negative control samples and were excluded from further analysis.

### Homology modelling, structure visualization and measuring cross-link distances

Homology models for the TBEV C protein, immature conformation of the E protein and the prM protein were generated using the I-TASSER (Iterative Threading ASSEmbly Refinement) server [35–37]. A C-score (confidence score for estimating the quality of predicted models by I-TASSER) is generated for each model and can range from -5 to 2, with a higher value signifies a higher confidence and where a C-score > -1.5 is considered good [35–37]. The C-protein homology model was generated using PDB accession 5OW2, and the immature conformation of the E protein, and the prM protein using PDB accession 7L30 as the template. The TBEV polyprotein sequence was obtained from GenBank (accession: AWC08512.1). The sequence for the C protein was obtained from residues 1-96, the E protein residue 281-776, and the prM residues 113-280 in the polyprotein. Generation of the assembled immature virus homology model, and the C-protein dimer was performed in UCSF Chimera [80]. The homology model for the immature virus was generated by superimposing the models for the immature E protein and prM protein onto the assembled immature Spondweni virus (PDB accession 6ZQW) structure, using the MatchMaker function. The homology model for the C protein dimer was generated by superimposing two models of the C protein onto the Zika C protein dimer (PDB accession 5YGH), using MatchMaker. The atomic models were used to position both intraprotein and interprotein cross-links by choosing the distance between Cα atoms in Chimera [25, 27].

### Networking and clustering analysis

String data were obtained from string database [41] and imported into Cytoscape 3.4 [81] .The following interaction sources were considered: experiments, databases, co-expression, co-occurrence and gene fusion, and interactions with a minimum interaction score of 0.7 are shown. Clusters were generated using the Markov clustering algorithm and gene ontology annotations for each cluster were obtained using GOnet [82].

### Data Availability Statement

The datasets generated during and/or analysed in the current study are available in the repositories with the persistent web links: ftp://massive.ucsd.edu/MSV000088272/ [76].

The mass spectrometry data has been deposited to the ProteomeXchange consortium via the MassIVE partner repository https://massive.ucsd.edu/ with the dataset identifier PXD029384 [76].

## Acknowledgments

We thank Markku Varjosalo and Johan Malmström for helpful discussions. The work was carried out in the Instruct Centre Finland supported by grants to SJB from the Swedish Research Council reference number 2018-05851; the Academy of Finland grant number 336471, the Sigrid Juselius Foundation grant number 95-7202-38. This project has also received funding from the European Union’s Horizon 2020 Research and Innovation Programme under the Marie Skłodowska-Curie grant agreements number 765042 (ViBRANT) and number 799929 (MA). SVB is a fellow of the Doctoral Programme in Integrative Life Science, University of Helsinki. LIAP is a fellow of the Doctoral Programme in Microbiology and Biotechnology, University of Helsinki. We gratefully acknowledge support from the Swedish National Infrastructure for Biological Mass Spectrometry.

## Conflict of interests

The authors declare that they have no conflict of interest.

## Supporting information captions

**S1 Table: DDA analysis of viral and host protein in all samples (XLSX).**

**S2 Table: Cross-linking dataset** Identified cross-linked peptides, corresponding proteins, E-values and SVM scores for each cross-linked spectral observation (XLSX).

**S3 Table: Filtered cross-linking dataset** Cross-linking dataset (S2 Table) filtered to remove spectral observations corresponding to unique cross-links identified by less than 2 spectral observations with SVM scores < 0.355 (XLSX).

**S4 Table: GO analysis of host proteins** GO analysis of the 59 host proteins shown to interact with the surface of TBEV in the filtered cross-linking dataset (S3 Table) (XLSX).

**S1 File: Correlation between number of cross-linked spectral observations, protein abundance and protein lysine content** Table showing the number of cross-linked spectral observations, the average protein abundance calculated in the DDA analysis (S1 Table) and the number of lysines for 101 randomly selected proteins. Scatter graphs show no correlation between the number of cross-linked spectral observations and the number of lysines for the proteins, and no correlation between the number of cross-linked spectral observations and the protein abundance (XLSX).

**S1 Fig: Box and whisker plot of the E-value for the intraviral cross-links, plotted against the distance between the cross-linked residues for each cross-link.** The smallest distance out of the mature and immature calculated distances, and the C protein dimer or monomer distances is plotted. The mean E-value is marked with a cross.

## Notes

ftp://massive.ucsd.edu/MSV000088272/

### Competing Interest Statement

The authors have declared no competing interest.

### Summary of Updates

Discussion revised Reference added Acknowledgements revised

