## Supplementary figures and images for "Proteome-wide cross-linking mass spectrometry to identify specific virus capsid-host interactions between tick-borne encephalitis virus and neuroblastoma cells"

### Supplemental Figure 1

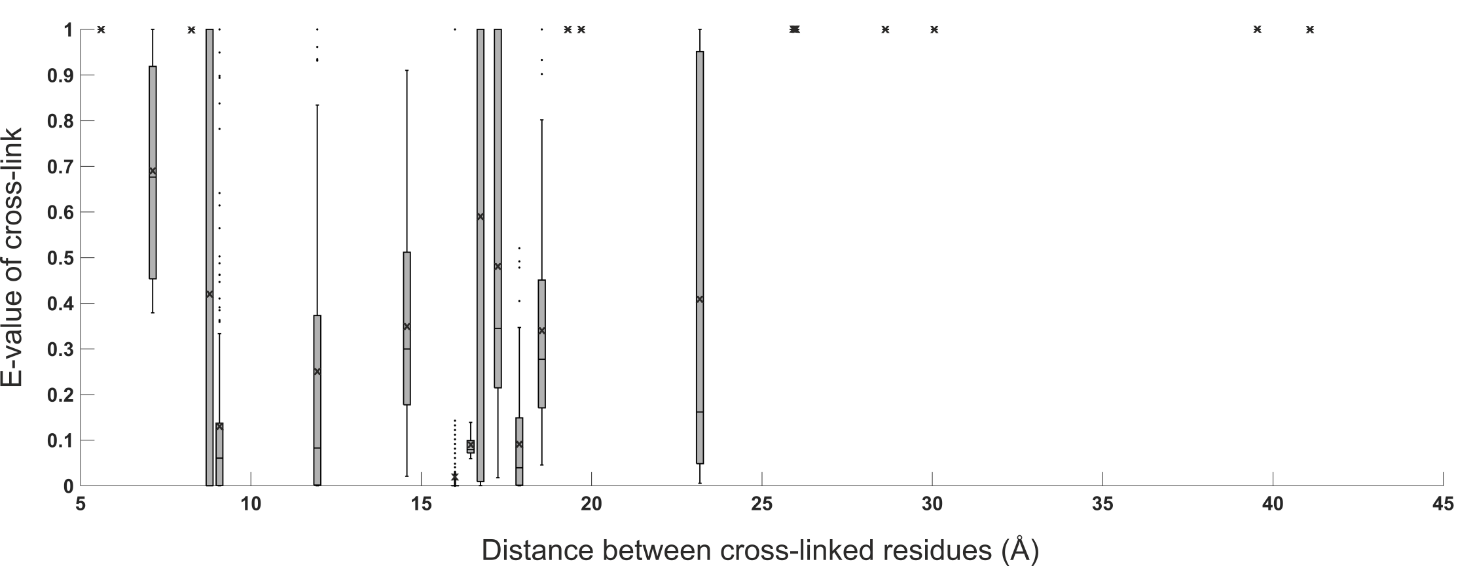
